# 3-Phosphoinositide-dependent kinase 1 drives acquired resistance to osimertinib

**DOI:** 10.1101/2021.12.10.472153

**Authors:** Ismail M. Meraz, Mourad Majidi, Bingliang Fang, Feng Meng, Lihui Gao, RuPing Shao, Renduo Song, Feng Li, Min Jin Ha, Qi Wang, Jing Wang, Elizabeth Shpall, Sung Yun Jung, Franziska Haderk, Philippe Gui, Jonathan Wesley Riess, Victor Olivas, Trever G Bivona, Jack A. Roth

## Abstract

Osimertinib sensitive and resistant NSCLC NCI-H1975 clones were used to model osimertinib acquired resistance in humanized mice and delineate potential resistance mechanisms. No new EGFR mutations or loss of the EGFR T790M mutation were found in resistant clones. Resistant tumors in humanized mice were initially partially responsive to osimertinib, then aggressive tumor regrowth occurred accompanied by an immunosuppressive tumor microenvironment. 3-phosphoinositide-dependent kinase 1 (PDK1) was identified as a potential driver of osimertinib acquired resistance, and its selective inhibition by BX795 and CRISPR gene knock out, sensitized resistant clones and a patient derived xenograft (PDX) with acquired resistance to osimertinib. PDK1 knock-out dysregulated PI3K/Akt/mTOR signaling, promoted cell cycle arrest at the G1 phase, and inhibited nuclear translocation of yes-associated protein (YAP). Higher expression of PDK1 was found in patients with progressive disease following osimertinib treatment. PDK1 is a central upstream regulator of two critical drug resistance pathways: PI3K/AKT/mTOR and YAP.

## Introduction

Tyrosine kinase inhibitors (TKIs) targeting the epidermal growth factor receptor (EGFR) have become the standard of care for NSCLC patients with EGFR driver mutations^1,2^. Osimertinib (AZD9291) is the first FDA-approved third-generation EGFR TKI, which irreversibly binds to EGFR proteins with T790M drug resistance mutations^3–6^. Not all patients respond initially, and responses, when they occur, are variable, typically incomplete, and temporary due to acquired drug resistance^7–15^. This obstacle to long-term patient survival highlights the need to identify acquired resistance mechanisms. Acquired drug resistance is a complex problem as multiple downstream effector proteins in bypass pathways can drive tumor regrowth, progression, and ultimately drug resistance^16–19^.

The PI3K/AKT/mTOR has been implicated in NSCLC acquired resistance, but the role of AKT-independent signaling downstream of PI3K is not well-characterized^20,21^. One protein that transduces PI3K signaling, is the serine/threonine kinase 3-phosphoinositide-dependent protein kinase 1 (PDK1 also known as PDPK1). PDK1 is a pleotropic regulator of 60 serine/threonine kinase proteins classified into 14 families of the AGC kinase superfamily^22^. It has multifunctional oncogenic activity, concurrently activating pro-survival protein kinases^23,24^, and suppressing apoptosis in lung cancer^25^. PDK1 is also implicated in tumors driven by KRAS genetic alterations, and regulates immune cell development, including T and B cells, and their functions in the tumor microenvironment (TME)^26–29^. NSCLC sera compared to healthy samples were reported to have significantly higher levels of PDK1 mRNA expression^30^. Recently, PDK1 started to gain a wide interest as a drug target, which has so far led to the filing of more than 50 patents^24^.

The validity of current preclinical modeling of osimertinib acquired resistance has been questioned because of inaccuracy in predicting clinical benefit. Studies have relied on mouse tumor biology, which is useful to certain degree, but cannot provide an accurate recapitulation of osimertinib interactions with a highly complex and heterogeneous human tumor microenvironment (TME). There is still a need for a reliable preclinical platform to model osimertinib acquired resistance. We have recently published a humanized mouse model using irradiated NOD.Cg-*Prkdc^scid^ Il2rg^tm1Wjl^*/SzJ (NSG) mice injected with fresh, non-cultured, human CD34^+^ stem cells with rapid and prolonged reconstitution of multiple functional human immune cell subpopulations. This model a) replicates human tumor response to checkpoint blockade, b) mounts an immune response to tumor-associated antigens and not alloantigens, and c) is not stem cell donor–dependent for its immune response^31^.

In this study, we used this humanized mouse model implanted with the human NSCLC osimertinib sensitive xenograft, NCI-H1975 harboring EGFR T790M and L858R point mutations, and its osimertinib resistant isogenic xenograft, NCI-H1975/OSIR, to model osimertinib acquired resistance. We provide evidence that a) PDK1 is a potential driver of osimertinib acquired resistance, b) PDK1 genetic and pharmacological targeting restores osimertinib response in resistant clones and their derived human xenografts and PDXs, c) immune contextures of osimertinib sensitive and resistant humanized xenografts have immunologically distinct TMEs, and d) patient biopsies from EGFR mutant lung adenocarcinoma tumors with the highest PDK1 expression associate with patient’s progressive disease following TKI treatment, suggesting they could be responsive to PDK1 inhibitors.

## Materials and Methods

### Osimertinib resistant cells, cell culture and maintenance

The human parental NCI-H1975 NSCLC cell line, which carries EGFR T790M/L858R mutations, and its osimertinib resistant isogenic clone, NCI-H1975/OSIR, were obtained from Dr. John Minna’s laboratory (University of Texas Southwestern University, Dallas, TX), and Dr. John Heymach’s laboratory (University of Texas MD Anderson Cancer Center (MDACC), Houston, TX), respectively. NCI-H1975/OSIR cells were cultured and maintained in RPMI complete media supplemented with 1 µM osimertinib (Medchemexpress (MCE), NJ, USA), which was dissolved in DMSO, stored at -70°C, and diluted in culture medium for in-vitro experiments.

### Antibodies

Antibodies were purchased from Cell Signaling (Beverly, MA): anti-AKT (CS#4691), p-AKT (Ser473) (CS#9271), p-AKT (Thr308) (CS #4056), mTOR, p-mTOR (Ser2448) (CS#2971), PDK1 (CS#13037), p-PDK1(Tyr373/376) (bs-3017R), p-PDK1(Ser241) (CS#3061), MAPK (CS#4695), p-MAPK (Thr202/Tyr204) (CS#4370), PTEN (CS#9559), p-PTEN (CS#9549)). Monoclonal anti-β-actin (Sigma#A5449) was purchased from Sigma Aldrich (St Louis, MO).

### Generation of PDK1-knockout and PDK1 overexpression cells

NCI-H1975/OSIR PDK1-knockout clones were generated with CRISPR-Cas9 technology (CRISPR core lab, Baylor College of Medicine (BCM), Houston, TX). The PDK1 overexpressing clone was generated by the BCM core lab, by stable transfection with Myc-tagged PDK1 overexpressing plasmid (OriGene, Rockville, MD).

### Whole Exome Sequence Analysis

NCI-H1975 and NCI-H1975/OSIR cells were seeded in triplicates at a cell density of 2 X 10^6^/plate. DNA was isolated and purified using a Qiagen kit (Germantown, MD). The quality of DNA was evaluated and the whole exome sequenced at the sequencing core lab, MDACC, Houston, TX using a next generation sequencer (NextSeq500, Illumina, USA). Sequencing data were analyzed by the Department of Bioinformatics and Computational Biology, MDACC.

### Drug sensitivity assay

NCI-H1975 and NCI-H1975/OSIR isogeneic cells were seeded at a density of 3X10^3^ cells/well in a 96-well microplate and treated with osimertinib at concentrations ranging from 0.001 to 10 µM in DMSO. Cells were incubated in 37°C incubators, 5% CO_2_, for three days. Cytotoxicity assays were performed using colorimetric XTT (Sigma-Aldrich, USA) and SRB (Sulforhodamine B) Assay Kit (Abcam, USA) reagents according to manufactured protocol. Optical density (OD) was measured using a microplate reader (FLUOstar Omega, BMG Labtech USA) at 570 nm. Cell viability (%) = [OD (Drug) - OD (Blank)] / [OD (Control) - OD (Blank)] × 100.

### Development of osimertinib sensitive and resistant tumor xenografts in humanized mice

All animal experiments were carried out following approval by the MDACC institutional review board and were performed in accordance with the Guidelines for the Care and Use of Laboratory Animals published by the National Institutes of Health. All measurements quantifying experimental outcomes were blinded to the intervention. After mononuclear cells were separated from human umbilical cord blood, CD34^+^ HSPCs were isolated using a CD34^+^ MicroBead kit (Miltenyi Biotec). Three- to 4-week-old NSG mice were irradiated with 200 cGy using a ^137^Cs gamma irradiator. Over 90% pure freshly isolated HLA typed CD34^+^ HSPCs were injected intravenously, twenty-four hours post irradiation, at a density of 1 to 2 × 10^5^ CD34^+^ cells/mouse. Ten mice per group from multiple umbilical cord blood donors were used. The engraftment levels of human CD45^+^ cells were determined in the peripheral blood, as early as 4 weeks post CD34 injection, by flow cytometric quantification, as well as other human immune populations. Mice with 25% human CD45^+^ cells were considered as humanized (Hu-NSG mice). The reconstitution levels of human immune cell populations in mice was analyzed throughout experiments using a 10-color flow cytometry panel at week 6 post CD34^+^ engraftment. These include CD45^+^, CD3^+^, CD4^+^ and CD8^+^ T cells, B cells, NK cells, dendritic cells (DC), myeloid derived suppressor cells (MDSC), and macrophages. Hu-NSG mice from different cord blood donors with different levels of engraftment were randomized into every treatment group in every experiment. All Hu-NSG mice were verified for humanization before tumor implantation. Treatment strategies for different experiments are described in the figures.

## Generation of PDXs with acquired resistance to osimertinib

To develop NSCLC PDXs with acquired resistance for osimertinib, we monitored mice with regressed tumors for tumor re-growth. When those tumors regrew to 200 mm^3^ in size, we retreated, until mice were euthanized. The tumors were passaged to new NSG mice for osimertinib sensitivity testing. PDX TC386, is in passage 4, with each generation treated with three or more cycles. Susceptibility to osimertinib was reduced in each passage. We performed whole exome sequencing for two tumors obtained in passage 3 (G3) of TC386 that was under constant osimertinib treatment and became less responsive. Both G3R1 and G3R2 had the same EGFR exon 19 deletion as the primary tumor (TC386T) and parental PDX (TC386F2), albeit with increased allele frequencies without novel EGFR mutations including absence of T790M. The PDXs with acquired resistance had new mutations that were not detected in either the primary tumor or parental PDX, including mutations in FAT3 and SETD1B.

### Immune profile analysis by flow cytometry

Harvested fresh tumors were processed for single-cell suspensions by enzymatic digestion (Liberase Enzyme Blend, Roche, USA). Erythrocytes in the peripheral blood were lysed with ACK lysis buffer (Fisher Scientific). Several 10-color flow cytometry panels were used for immune profiling of innate and adaptive immune populations. Fluorochrome–conjugated monoclonal antibodies to the following human antigens were used: CD45-Alexa Fluor 700 (clone 2D1, HI30), CD45-phycoerythrin (PE; clone 2D1, HI30), CD3-PerCp/cy5.5 (clone HIT3a), CD19-PE-cyanine 7 (clone HIB19), CD8-allophycocyanin-cyanine 7 (clone RPA-T8, HIT8a), CD4-Pacific blue (clone OKT4), CD56-PE/BV510 (clone HCD56), CD69-FITC/APC/PE-Alexa Flour 610 (clone FN50; Thermo fisher), HLA-DR-PerCp/cy5.5 (clone LN3), CD33-PE (clone WM-53) (Thermo fisher), CD11b-PE-Cy7 (clone 1CRF-44) (Thermo fisher), Granzyme B-FITC (clone GB11), and IFN-γ-APC (clone 4S.B3), CD103-Super bright 600 (Colne B-LY7; Thermo fisher), CD279 (PD-1)-Super Bright 702 (Clone J105; Thermo fisher), CCR7-FITC (Clone G043H7), CD45RA-PE (Clone HI100), CD25-APC (clone CD25-4E3), Lin-FITC (Biolegend), CD163-APC (clone ebioGH1/61; Thermo fisher), CD11c-Pacific blue (clone Bu15; Thermo fisher). A mouse CD45-FITC (clone 30-F11) antibody was used for gating out murine leukocytes. Antibodies were purchased from Biolegend. All samples were run on Attune NxT flow cytometer (Thermo fisher), and data were analyzed by Flow Jo software.

### Development of osimertinib sensitive and resistant tumor xenografts in non-humanized mice

NCI-H1975/OSIR isogenic cells were cultured and expanded in osimertinib (1uM) containing media. NCI-H1975 and NCI-H1975/OSIR at 5X10^6^ cell density were injected subcutaneously into 6-8 weeks old NSG mice. When tumor size reached approximately 100 mm^3^, tumor-bearing mice were randomized and treated with osimertinib, 5mg/kg or 10mg/kg, 5 days a week for 3 weeks. To evaluate the effect of the PDK1 selective inhibitor, BX795, 5 tumor bearing mice/group were either left untreated, treated with osimertinib alone (5mg/kg), treated with PDK1 inhibitor BX 795 (25mg/kg) (Selleckchem, Houston, TX), or with the combination. Cells (5X10^6^) were injected subcutaneously into 6-8 week old NSG mice followed by osimertinib treatment starting from the day following tumor cell injection so tumors developed under osimertinib pressure. Mice were treated with osimertinib from the first day of tumor cell injection throughout the entire experiment, and tumor sizes were measured twice a week by caliper. Tumor volume was measured using the formula V=ab^2^/2 where a is the largest diameter and b is the smallest. At end of the experiment, residual tumor tissues were harvested for further analysis.

### In-vivo inhibition of PDK1 in an osimertinib resistant NSCLC PDX

To evaluate the effect of PDKi, BX 795, we propagated the EGFR mutant TC386R PDX with acquired osimertinib resistance in NSG mice. After 3 weeks, fresh PDXs were harvested and 2X2 cm size PDX tissues were re-implanted into 25 NSG mice for the experiment. Large size PDX bearing mice were randomized into four groups including control, osimertinib alone (10mg/kg), BX 795 alone (25mg/kg) and an osimertinib + BX 795 combination group. Osimertinib treatment was 5 days a week (oral) and BX 795 treatment was 2 times per week (i.p.). Tumor size was measured twice a week by caliper. Tumor volume was calculated according to the formula V=ab^2^/2, where a is the largest diameter and b is the smallest. At the end of the experiment, residual tumor tissues were harvested for analysis.

### Western blot analysis

Total protein was harvested using Ripa lysis buffer (Merck, Burlington, MA), and their concentrations were evaluated with BCA^TM^ protein assay kit (Pierce, Rockford, IL, USA). Equal amounts of proteins were separated by 8-15% SDS-PAGE gel, electro-transferred onto a Hybond ECL transfer membrane (Amersham Pharmacia, Piscataway, NJ), and blocked with 2-5% non-fat skim milk. Then, membranes were probed with specific primary antibodies overnight at 4 °C, washed with PBS, and incubated with corresponding secondary antibodies at room temperature for 1 h. The specific protein bands were visualized with an ECL advanced western blot analysis detection kit (GE Health Care Bio-Sciences, NJ, USA).

### Colony formation assay

NCI-H1975, NCI-H1975/OSIR, NCI-H1975/OSIR-PDK1-/- and NCI-H1975-OSIR/PDK1++/++ cells were seeded into 6-well plates at a density of 300 cells per well, and treated with 0, 10nM, 100nM, 1 µM, 2.5 µM, and 5 µM concentrations of osimertinib for 72 h. The media was replaced every 2-3 days with osimertinib dose titration containing medium. After twelve days, colonies were fixed with 4% PFA, stained with crystal violet solution, and photographed.

### Cell Cycle assay

The cell cycle profiles of osimertinib resistant and sensitive cells were determined by staining DNA with fluorescent dye (PI/RNase staining buffer, BD Pharmingen, USA) according to the manufacturer’s protocol, and measuring its intensity by flow cytometry (Attune NxT, Thermofisher Scientific, USA). Briefly, cells were seeded at 10^6^ cells in a 100mm dish followed by osimertinib treatment at the designated concentrations. Cell pellets were suspended in ice cold PBS, fixed with 70-80% ethanol, and stored at -20^0^C overnight. The cells were washed twice with ice cold PBS and stained with PI/RNase staining dye for 15 min at room temperature. The samples were analyzed by flow cytometry within an hour.

### Reverse Phase Protein Array (RPPA) analysis

Osimertinib sensitive and resistant H1975 tumors were developed in non-humanized NSG mice and treated with osimertinib under the protocol described above. Residual tumor tissues were snap-frozen and stored in -80^0^C. RPPA analysis was performed using 400 antibodies at the RPPA core lab at MDACC. Bioinformatics analysis was performed by the Department of Bioinformatics and Computational Biology, MDACC.

### Mass Spectrometry

The samples were denatured and lysed by three cycles of LN2 snap freeze and thaw at 95°C. For global profiling, 10 mg of lysate was trypsinized to obtain 10 mg of digested peptides. After fractionation using a small-scale basic pH reverse phase (sBPRP) step elution protocol with increasing acetonitrile concentrations, fractions were combined into 5 pools that were resolved and sequenced online with Fusion Lumos and timsTOF fleX mass spectrometer. For phospho-proteome profiling, a 100 µg protein lysate was digested with trypsin and dried under vacuum. Global and phospho-proteomic analyses covered over 8000 gene protein products (GPS), and over 4000 GPS, respectively, which included the kinome profile. After label-free nanoscale liquid chromatography coupled to tandem mass spectrometry (nanoLC-MS/MS) analysis using Thermo Fusion Mass spectrometer, the data was processed and quantified against NCBI RefSeq protein databases in Proteome Discover 2.5 interface with Mascot search engine (Saltzman, Ruprecht). The Skyline program was used to get precise quantification. To decipher phospho-proteome signal pathway analysis we utilized protein external data contributions for phosphorylation-related data mining sets, including PhosphoSitePlus (http://www.phosphosite.org/), Phospho.ELM, PhosphoPep, and the Phosphorylation Site Database (PHOSIDA).

### Immunofluorescence

Cells were seeded at 5000 cells/chamber well and grown overnight before being treated with osimertinib at 1uM or 2.5uM for 24 hours. Then cells were washed with PBS and fixed in 4% paraformaldehyde in PBS pH7.4 for 10 min at RT. After washing with PBS 3 times, cells were treated with 0.125% Triton-X100 for 10 min at RT to increase cell permeability. The slides were blocked by 1%BSA block in PBS-T (Thermo-Fisher) at RT for 30min, incubated with 1:250 anti-pYAP Y357 (Sigma #Y4646) antibody in 1%BSA overnight at 4C. After three PBS-T washes, the slides were further incubated in 1:1000 secondary Alexa 594 antibody (invetrogen #A32741) at RT for 1 hour, before they were mounted with mounting media containing Dapi (abcam #104139). Immune fluorescence images were captured using an EVOS M5000 fluorescence microscope (Thermos-Fisher). For each cell line and each treatment condition, 30 individual cell nuclei were counted and their fluorescent intensities were quantified.

### Immunohistochemistry

Clinical specimens of NSCLC samples that were obtained from patients before initiating systemic targeted therapy (TKI naive [TN]), at the residual disease (RD) state, and upon subsequent progressive disease as determined by clinical imaging, at which point the tumors showed acquired drug resistance (progression [PD]). All patients gave informed consent for collection of clinical correlates, tissue collection, research testing under Institutional Review Board (IRB)-approved protocols. Patient studies were conducted according to the Declaration of Helsinki, the Belmont Report, and the U.S. Common Rule. Formalin-fixed paraffin-embedded (FFPE) tumor blocks were cut at 4-micron thickness and mounted as sections on positively charged histology slides. Immunohistochemistry staining was performed as described previously. In brief, slides were deparaffinized in xylene, rehydrated and epitope retrieval was induced in a histology pressure cooker using pH 6.1 citrate buffer (Dako Denmark A/S, S2369). After endogenous peroxidase, tissue was permeabilized in in 0.1% Triton-X/PBS. Non-specific binding was blocked, and slides were incubated primary antibody solution overnight at 4 °C. The antibody for PDPK1 (clone EP569Y, # ab52893) was purchased from Abcam and diluted 1:150. Then, slides were incubated with secondary antibody for 30 minutes (EnVision Dual Link Labelled Polymer HRP, Agilent K4065), stained using 3,3-DAB, and counterstained with hematoxylin. Slides were dehydrated and mounted before digitization using an Aperio AT2 Slide Scanner (Leica Biosystems) at a 20X objective.

## Statistical analysis

### WES

The quality of raw FASTQ reads was assessed using FastQC and then mapped to human reference genome GRCh38, using BWA^32,33^. The reference genome refers to the b38 version with decoy sequences for human GRCh38 provided in the genome analysis toolkit (GATK) resource bundle^34^. The mutations were called following GATK best practice pipeline. The candidate mutations were be filtered for high confident somatic mutations and annotated for functional changes using ANNOVAR^35^.

### Cell Survival Assay

The percentage of viable cells was determined by the ratio of absorbance of treatment and control groups: ODT/ODC x 100%. Univariate analysis was performed to evaluate the distribution of data for each treatment group. To determine whether SRB % was different between treatment groups, two methods were used: 1) ANOVA was performed to compare the variance between treatment groups for all samples within each cell line; and 2) Tukey’s multiple comparisons test was performed for pairwise differences between treatment groups. P<0.05 was considered statistically significant; all tests were two-sided. Analyses were performed using SAS 9.3 (SAS Institute Inc., Cary, NC). Values represent the mean of three independent experiments.

### Colony Formation Assay

Colonies were fixed with glutaraldehyde, stained with crystal violet, counted with a stereomicroscope, and analyzed with Image-J software. Values represent the mean of three independent experiments. The statistical significance of differences between treatment groups was calculated by two-tailed t test analysis; P< 0.05 was considered significant. The Statistical software S-PLUS 8.0 was used for all analyses.

### Tumor growth

Statistical analyses were performed with GraphPad Prism 7 software. Tumor intensity change per time point was calculated as a relative level of tumor intensity change from baseline. Two-way ANOVA with interaction of treatment group and time point was performed to compare the difference of tumor intensity changes from baseline between each pair of the treatment groups at each time point. Means ± standard errors of the mean are shown in all graphs. The nonparametric Mann-Whitney U test was applied to compare cell numbers in different treatment groups. Differences of *P* < 0.05, *P* < 0.01, and *P* < 0.001 were considered statistically significant. Statistical analysis of flow cytometry data was done by general linear regression models to compare the data among the different treatment groups. CONTRAST statement in PROC GENMOD procedure in SAS was used to compare the data between each pair of the treatment groups with treatment indicator in the models. Both nom*P* values and multiple testing adjusted *P* values were reported. SAS version 9.2 and S-Plus version 8.04 were used for the computations for all analyses.

### Reverse Phase Protein Array (RPPA)

Slides were scanned using a CanoScan 9000F and spot intensities were quantified using ArrayPro Analyzer 6.3 (Media Cybernetics Washington DC). SuperCurve, a software developed in house, was used to estimate relative protein levels^36^.

After SuperCurve fitting, protein measurements were normalized for loading using median centering across antibodies. One-way analysis of variance (ANOVA) was used to assess the differences in protein expressions between control and treatment groups on a protein-by-protein basis. An over-all F test was carried out to detect any significant difference among the means of all groups. Fold change (FC) values between the 2 groups, with the following conventional modification: ratios > 1 (up-regulation), ratios ≤ 1 (down-regulation). The type I error rate of multiple comparison will be controlled by using the false discovery rate (FDR). The criteria of significant protein selection were: 1. Significant in overall F-test (FDR <0.05); 2. Significant in pairwise comparison (FDR <0.05).

### In-vivo inhibition of PDK1 in an osimertinib resistant PDX

We evaluated the potential synergistic effect of the drug combination osimertinib and BX795 under the Highest Single Agent framework^37^, where the synergistic effect of a drug combination is declared if the combination effect is greater than that of the more effective individual component. The combination index (CI) under the HSA and the corresponding standard error were approximated by the Delta method^38^. We declared the synergistic effect under the significance level of 5% at day 33.

### Mass Spectrometry

Statistical analysis was performed using R software (R version 4.0.1). The log2 transformation was applied to the iFOT Half Min proteomic data. The Student’s t-test was used to compare expression values between the groups. P values obtained from multiple tests were adjusted using FDR. Statistical significance was defined as FDR < 0.05. The enriched pathways and hallmarks were identified by pre-ranked GSEA using the gene list ranked by log-transformed P values with signs set to positive/negative for a fold change of >1 or <1, respectively.

### Immunohistochemistry (IHC)

To investigate the relations of YAP expression and PDPK1 groups, and PDPK1 %Cases and TN/PD, we used a Bayesian hypothesis testing framework to test whether samples from a category in columns are enriched in a category in rows. We compute the enrichment probabilities (EP) for each cell of the tables to represent the probabilities of patients in a column category are grouped together according to the row variable. We assume that the frequencies in each row in a table follow independent multinomial distributions, where the proportions have the prior distribution of Dirichlet (1,1,1,1). For a given category in columns, e.g., YAP=0, we compute the posterior probability that the samples with YAP=0 are clustered together in the PDPK1 LOW category rather than the High category, which constitutes the PDPK1 LOW and YAP=0 cell in the EP table.

### Immunofluorescence and others

we performed multiple t test analysis using GraphPad Prism 9 software.

## Results

### Osimertinib sensitivity assay

The human NSCLC NCI-H1975 cell line harbors two EGFR point mutations, T790M and L858R, in exons 20 and 21, respectively and is highly sensitive to osimertinib^39^. NCI-H1975 was continuously exposed to osimertinib dose escalation (0.5 µM -2.5 µM) until the emergence of the osimertinib resistant clone, NCI-H1975/OSIR, which has less longitudinal cell morphology and a faster doubling time than the parental sensitive clone (33hrs vs 42hrs). An osimertinib sensitivity assay using Sulforhodamine B (SRB) to measure cell proliferation showed that NCI-H1975/OSIR cells were 100-200 fold more resistant to osimertinib than their NCI-H1975 counterparts, as shown by IC 20, 30 and 50 values (Fig 1).

**Fig 1.**
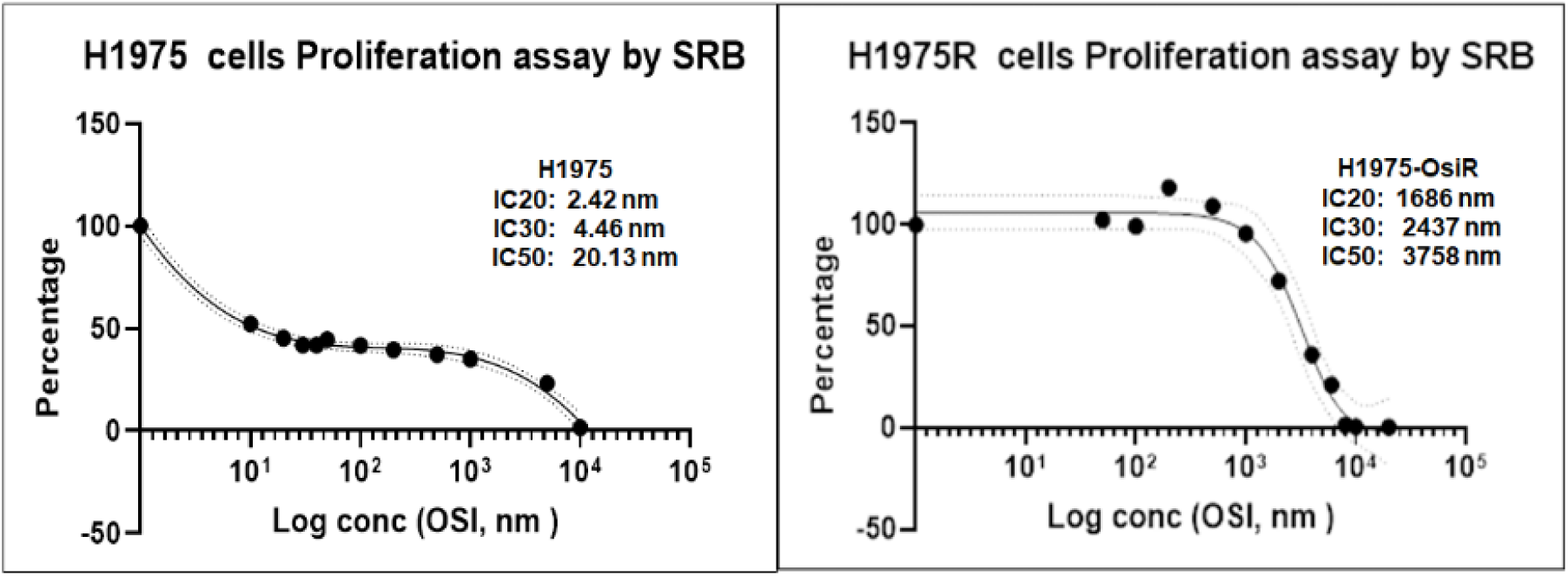
Effect of osimertinib on survival of NCI-H1975 and NCI-H1975/OSIR cells shown by IC 20, 30 and 50 values using the SRB assay. Data shown represent the mean ± SE of three independent experiments.

### Whole Exome Sequence (WES) analysis shows there are no new EGFR mutations or loss of the EGFR L858R and T790M mutations in NCI-H1975/OSIR

WES analysis was performed to identify any acquired or lost EGFR mutation during osimertinib treatment in the entire protein-coding regions of the genome of NCI-H1975/OSIR clones. The results showed there were no new EGFR mutations or loss of the EGFR L858R and T790M mutations. However, there were 37 new exonic mutations, including 0 indels, 27 nonsynonymous single nucleotide variants (SNV), 4 stopgain, and 6 synonymous mutations. The list of these new mutations and their projected pathways are shown in supplemental Fig 1.

### Modeling osimertinib acquired resistance in the humanized mouse model

The major limitation of current experimental rodent models is that many functional aspects of human innate and adaptive immunity cannot be recapitulated with mouse models. Our improved humanized mouse model is better suited to model osimertinib acquired resistance and provides insight into the complex interaction of osimertinib with variable contextures of the tumor microenvironment (TME).^31^ We found, as have others, that anti-tumor responses are independent of HLA status in humanized mice. In addition, when we used HLA-matched human bronchial endothelial cells in co-culture experiments, no allogenic responses were observed. NCI-H1975 and NCI-H1975/OSIR xenografts were implanted with fresh CD34^+^-derived humanized mice from different donors with partial HLA compatibility. The humanization protocol, the levels of human immune reconstitution in humanized mice, growth characterization of tumor xenografts and osimertinib treatment are illustrated in figure 2 (A-C). The level of human CD45^+^ and reconstituted T, B, and NK cells were high 78 days post CD34 engraftment. The general standard for mouse humanization is a minimum level of 25% of reconstituted human CD45 cells. Our improved humanization protocol achieved almost two-fold that level in less than six weeks. Results from two independent experiments with long (78 days) and short term (27 days) osimertinib treatment (5mg/kg), (Fig 2C and E), respectively, showed that growth of NCI-H1975 tumors were significantly inhibited. In contrast, NCI-H1975/OSIR treated with osimertinib showed initial growth stabilization followed, after a short time, by tumor regrowth. Because NCI-H1975/OSIR xenograft tumors were developed under constant osimertinib (5mg/kg) pressure, twenty-four hours post implantation, their growth was slower than their untreated NCI-H1975 counterparts. The osimertinib effect in this humanized mouse model was reproducible in multiple experiments (data not shown).

**Fig 2.**
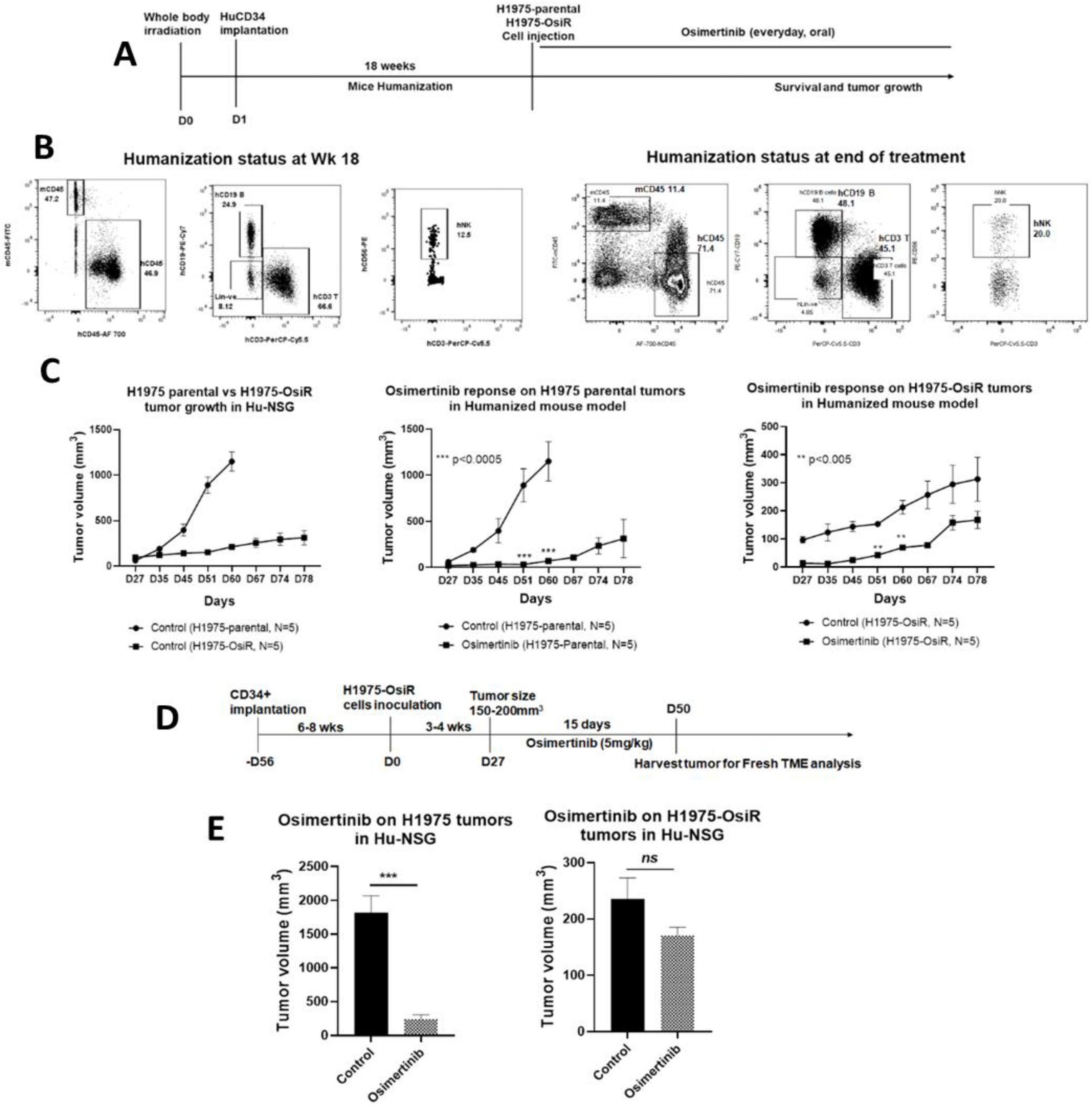
Effect of osimertinib on humanized H1975 and H1975/OSIR xenografts. **A)** Experimental strategy for mouse humanization, tumor cell inoculation, and osimertinib prolonged treatment, **B)** Levels of human immune cell repopulation in humanized mice at different time points. **C)** Tumor growth comparison between H1975-parental vs H1975-OsiR and the effect of osimertinib on their growth. **D)** Experimental strategy for mouse humanization, tumor cell inoculation, and osimertinib short term treatment. **E)** Antitumor effect of osimertinib on tumor growth.

### Analysis of the tumor microenvironment showed distinct immune contextures between osimertinib sensitive and resistant tumors

Fresh osimertinib sensitive and resistant tumor xenografts grown in humanized mice were processed into single cell suspensions and analyzed by multiple flow cytometry to identify differences in the immune contextures of both tumor microenvironments. Reconstituted human lymphoid and myeloid cell populations were investigated. Sensitive tumors had a higher number of CD11b^+^CD163^-^HLA-DR^+^ M1 macrophages than their resistant counterparts, which were infiltrated with a higher number of CD11b^+^CD163^+^HLA-DR^-^ M2 macrophages (Fig 3A). Osimertinib treatment significantly altered the ratio of M2 to M1, favoring the M1 population. The number of HLA-DR^+^ dendritic cells (DC) were higher in sensitive tumors, and osimertinib treatment enhanced their levels significantly, to the same extent in both sensitive and resistant tumors (Fig 3B). Surprisingly, CD33^+^ MDSC levels were higher in sensitive tumors, and were significantly reduced by osimertinib in both sensitive and resistant tumors (Fig 3C). The percentage of tumor infiltrating lymphocytes, CD3^+^TILs were moderately higher in sensitive tumors compared to their resistant counterparts and were significantly increased in both tumors following osimertinib treatment (Fig 3D). Natural Killer cells (CD56^+^ NK) were higher in resistant tumors and osimertinib enhanced their levels significantly more in resistant tumors (Fig 3E).

**Fig 3.**
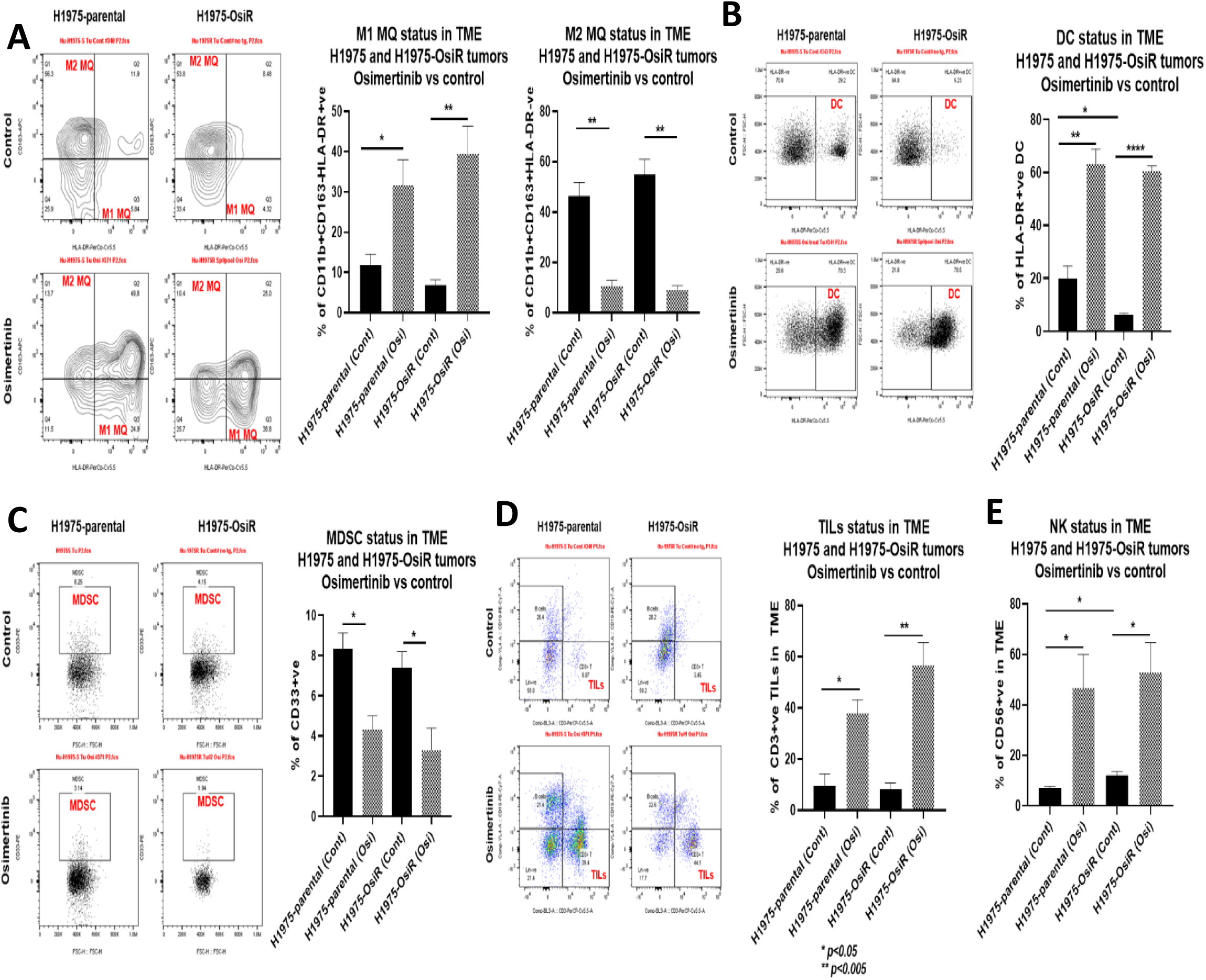
TME Immune profile analysis of humanized H1975 and H1975/OSIR tumors: **A)** human M1 and M2 macrophages, **B)** human dendritic cells, **C)** human MDSC, **D)** human tumor infiltrating lymphocytes (TIL) and **E)** human NK cells.

### Reverse Phase Protein Array (RPPA) shows upregulation of PDK1 in NCI-H1975/OSIR xenografts

To understand the underlying mechanisms of osimertinib acquired resistance, NCI-H1975 and NCI-H1975/OSIR xenografts were grown in NSG mice untreated or treated with 5 or 10mg/kg osimertinib. Tumors were grown under constant osimertinib pressure according to the treatment strategy shown in fig 4A. A dose dependent osimertinib response was observed in H1975-parental tumors (Fig 4B, D), which was completely lost in H1975-OSIR tumors (Fig 4C-D). Untreated and osimertinib treated residual tumors were harvested for proteomic analysis using an antibody-based functional proteomic analysis, RPPA, consisting of a 400 antibody panel, which includes serine/threonine and tyrosine kinases. Heat map analysis of pairwise comparison of protein expression profiles between NCI-H1975 and NCI-H1975/OSIR osimertinib treated residual tumors, and NCI-H1975/OSIR osimertinib treated and untreated groups showed upregulation of two distinct protein signatures, respectively. Interestingly in both signatures, PDK1 was a highly significant outlier and upregulation was highest in the osimertinib resistant tumors. In the former pairwise comparison, PDK1 expression level increased by 2.783 fold (Fig 4E), and in the latter by 2.4 fold log10 scale (Fig 4F). This suggests that PDK1 differential expression between NCI-H1975 and NCI-H1975/OSIR might play a potential role in the latter’s acquisition of resistance to osimertinib. PDK1 regulates a number of serine/threonine protein kinases of the AGC kinase superfamily, activating multiple pro-survival and oncogenic pathways, and suppressing apoptosis in lung cancer^22–25^.

**Fig 4.**
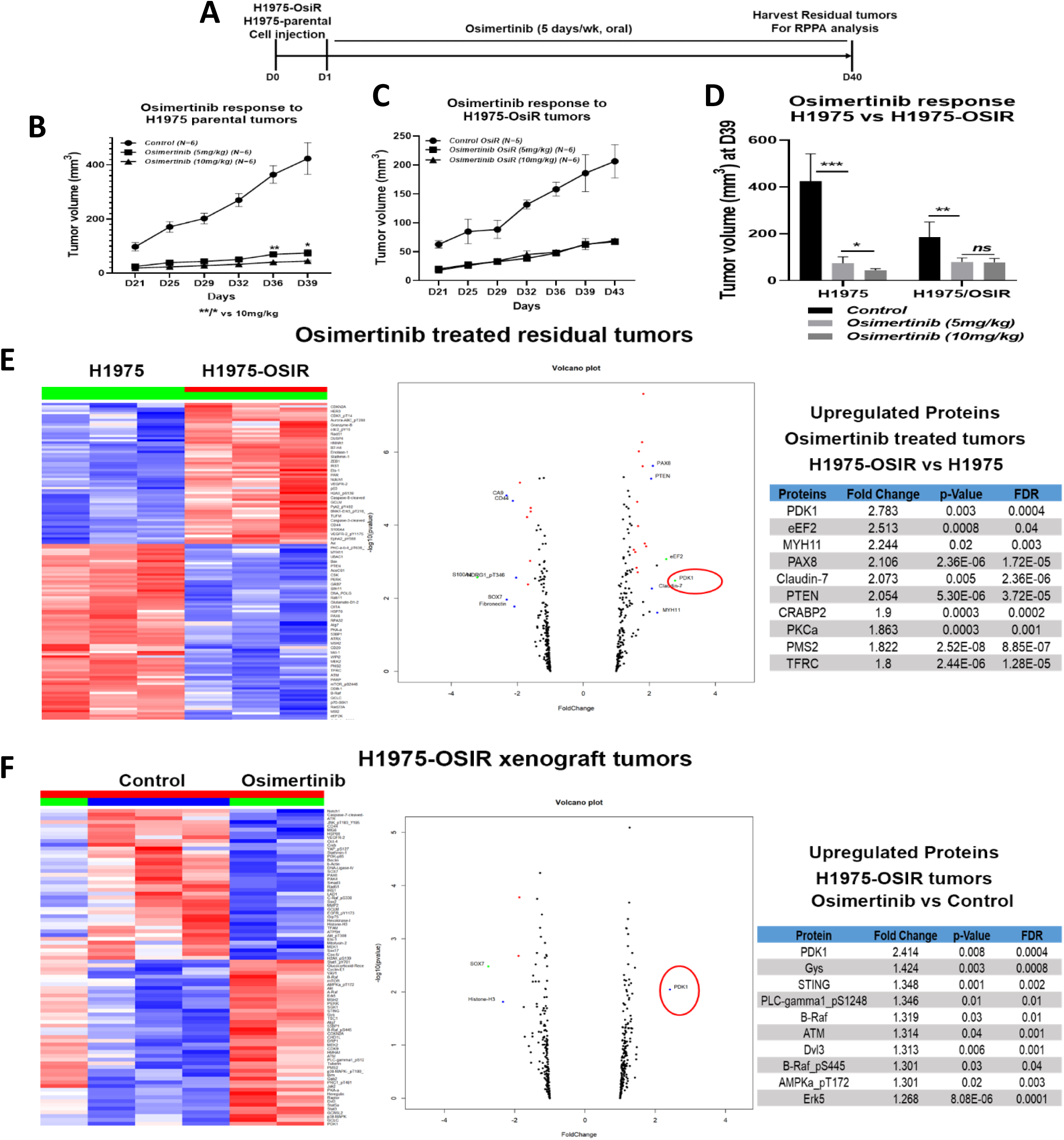
RPPA Gene expression profile analysis in osimertinib treated residual xenograft tumors. A) treatment strategy, B) osimertinib response to H1975 parental tumors, C) Osimertinib response to H1975-OSIR tumors, D) Osimertinib response at D39, E) Pairwise comparison between H1975 and H1975/OSIR residual tumors treated with osimertinib, F) Pairwise comparison between H1975-OSIR pre- and post-osimertinib treatment. The criteria of protein selection for each pairwise comparison were: 1. Significant in overall F-test (adjusted p value [FDR q values] <0.05); 2. Significant in pairwise comparison. (adjusted p value [FDR q values] <0.05). Heatmap/Volcano plot: proteins with FDR of 0.0001 and fold change >2.

### Mass spectrometry-based proteomic analysis confirms significant upregulation in PDK1 activity in osimertinib resistant NCI-H1975/OSIR clones

To further investigate the potential role of PDK1 in mediating osimertinib acquired resistance, as suggested by RPPA analysis, we profiled osimertinib proteins in the global and phospho-proteome of NCI-H1975 and NCI-H1975/OSIR isogenic clones using an unbiased robust mass spectrometry (MS)-based proteomics workflow^26,27^. Global and phospho-proteomic analyses covered over 8000 gene protein products (GPS), and over 4000 GPS, respectively. After label-free nanoscale liquid chromatography coupled to tandem mass spectrometry (nanoLC-MS/MS) analysis using a Thermo Fusion Mass spectrometer, the data was processed and quantified against NCBI RefSeq protein databases in a Proteome Discover 2.5 interface with Mascot search engine (Saltzman, Ruprecht). The level of PDK1 phosphorylation in NCI-H1975/OSIR, was fourteen fold higher than in the NCI-H1975 osimertinib sensitive cells (supplemental Fig 2). These results are compatible with those of RPPA, validating PDK1 differential expression and activity between NCI-H1975 and NCI-H1975/OSIR cells and identifying PDK1 as a potential driver of osimertinib acquired resistance.

### Pharmacological inhibition and genetic knock-out of PDK1 sensitizes NCI-H1975/OSIR clones to osimertinib

Next, we validated the functional role of PDK1 in osimertinib acquired resistance in vitro and in vivo. NCI-H1975/OSIR clones were left untreated, treated with osimertinib, the PDK1 selective inhibitor BX-795, or the combination of both, and assayed for survival by XTT assay. NCI-H1975 cells were used as controls. Figure 5A shows dose-dependent inhibition of pPDK1 expression by BX795 in both isogenic clones by western blot analysis. The osimertinib and BX-795 combination had no effect on the survival of NCI-H1975, whereas it rendered NCI-H1975/OSIR sensitive to osimertinib, as shown by a significant increase in cell death (Fig 5B). Next, to definitively implicate PDK1 as mediator of osimertinib acquired resistance and eliminate drug off-target effects, CRISPR PDK1 knockout (H1975-OsiR-PDK1-/-) and NCI-H1975/OSIR stably overexpressing PDK1 (H1975-OsiR-PDK1++/++) cells were generated (Fig 5C) and assayed for survival and colony formation following osimertinib treatment. PDK1 knockout sensitized NCI-H1975/OSIR to osimertinib, whereas PDK1 overexpression enhanced their resistance to osimertinib (Fig 5D). Similarly, in dose titration experiments, at nanomolar and micromolar osimertinib concentrations, PDK1 knockout and overexpression significantly reduced and enhanced NCI-H1975/OSIR colony formation, respectively (Fig 5E-F).

**Fig 5.**
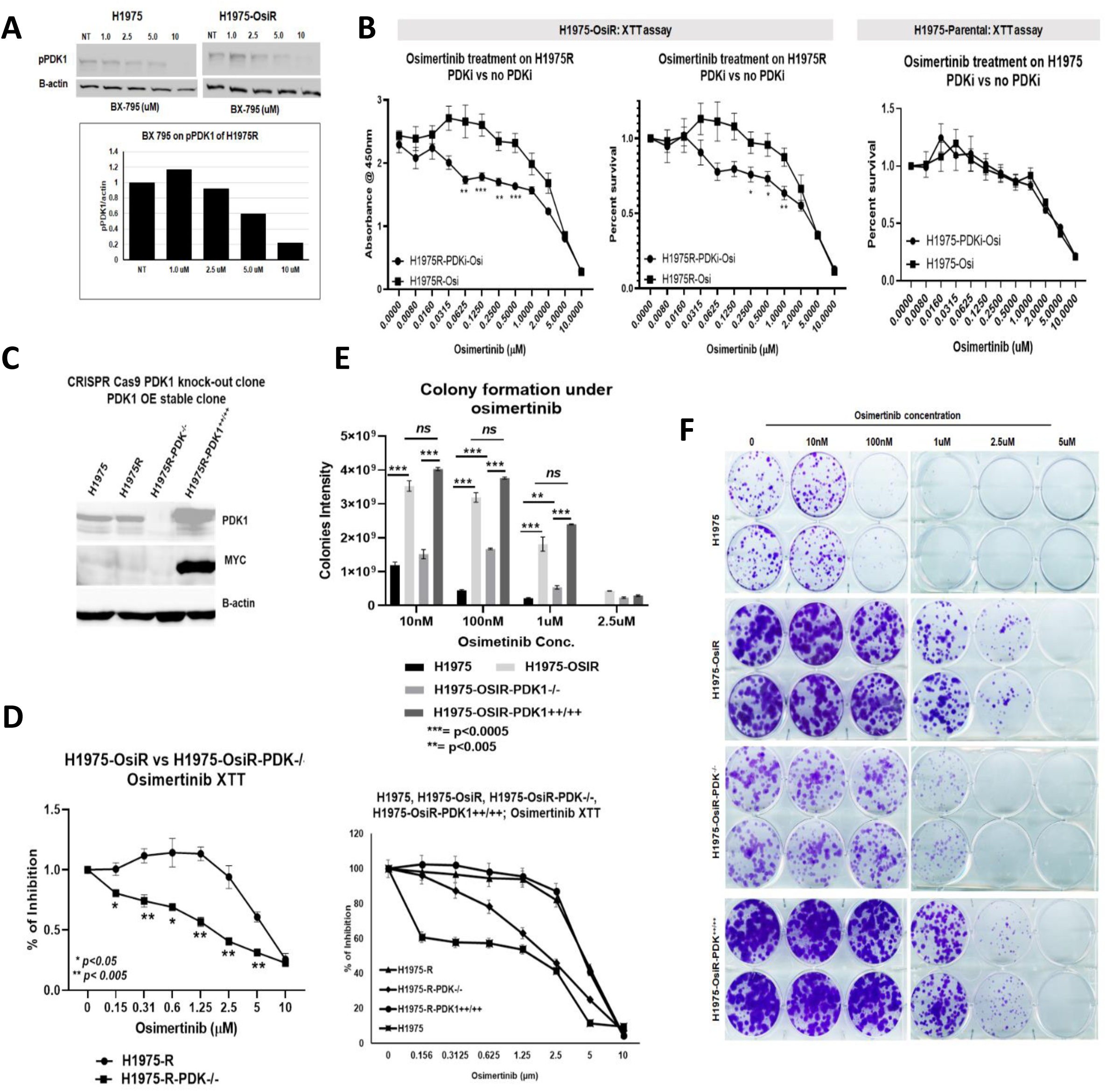
Effect of PDK1 knock out and overexpression on cell survival and colony formation. A) Dose dependent inhibition of pPDK1 by the PDK1 inhibitor, BX-795; B) XTT assay shows that BX795 renders NCI-H1975 OSIR sensitive to osimertinib; C) H1975 OSIR PDK1 knock-out and overexpressor clones; D) XTT assay shows the effect of PDK1 KO and overexpression on osimertinib response. E) Colony formation assay shows osimertinib differential sensitivity among all clones. Data shown represent the mean ± SE of three independent experiments.

### In-vivo inhibition of PDK1 enhanced osimertinib response in resistant PDXs

To evaluate the antitumor effect of PDK1 inhibition, we developed EGFR mutant osimertinib resistant TC386 isogenic PDXs. The parental TC386 PDX was highly sensitive to osimertinib (Supplemental Fig 3). The resistant TC386R PDX was generated through continuous treatment of osimertinib over a prolonged period of time and subsequent passages to four generations (Supplemental Fig 4). Later generation (RG4) showed significantly more resistance than an earlier generation (RG1) without the acquisition or loss of EGFR mutations identified in the parental PDX. When the pPDK1 level was compared between parental and resistant PDXs, higher levels of pPDK1 were found in TC386R PDXs as compared with parental TC386 PDXs (Fig 6A-B). To evaluate the effect of the PDK1 inhibitor (PDKi), BX 795 on resistant PDXs, we treated TC386R PDXs according to the treatment strategy shown in fig 6C treating with BX 795 with or without osimertinib. The combination treatment inhibited the tumor growth significantly and synergistically compared to the single agents (Fig 6D-E). BX 795 greatly reduced pPDK1 in the BX 795 tumors compared to untreated or osimertinib alone treated tumors (Fig 6A). Taken together, the in vitro and in vivo evidence support PDK1 as a driver of osimertinib acquired resistance in two independent models.

**Fig 6.**
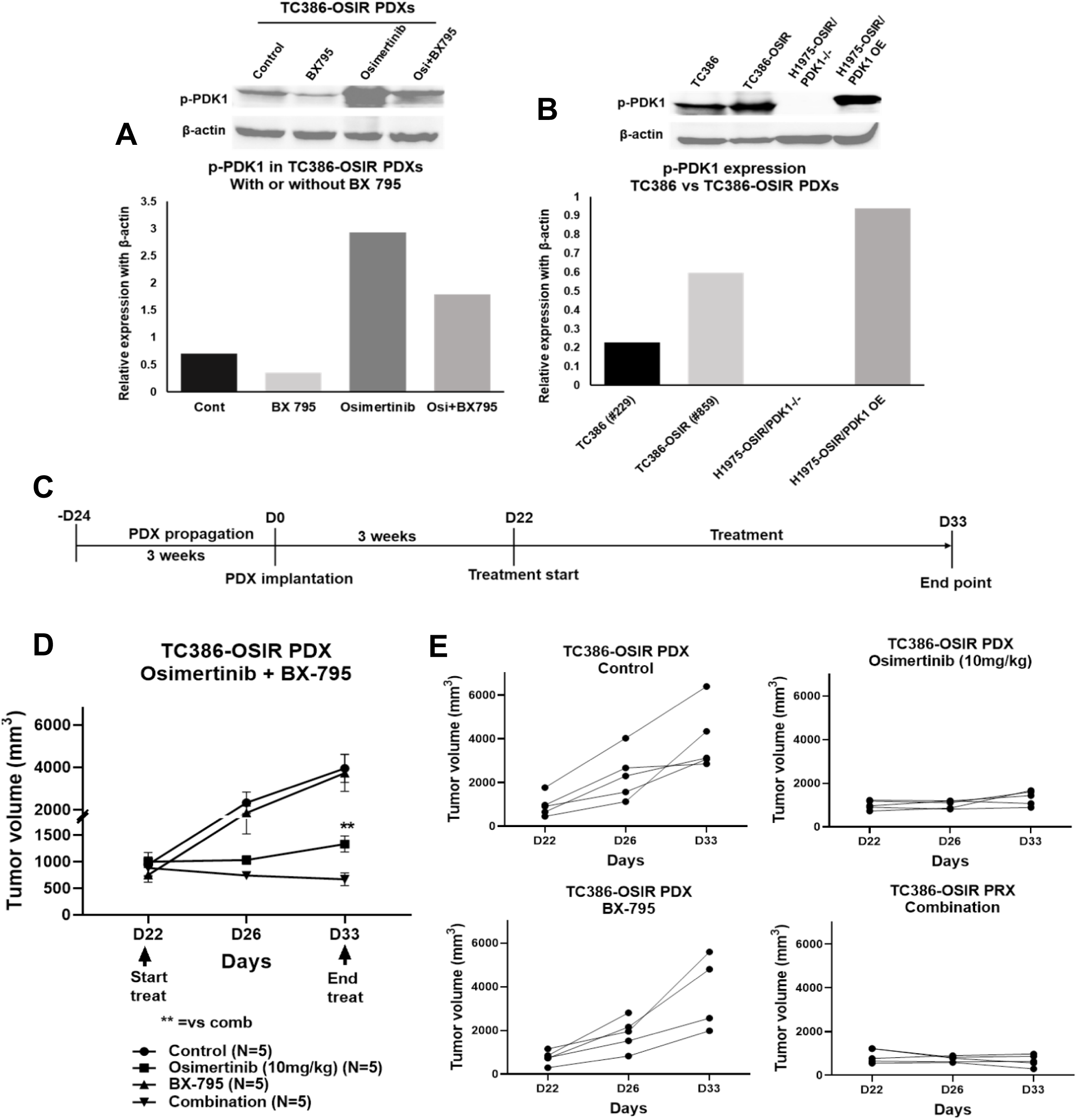
In-vivo inhibition of PDK1 by PDK1 inhibitor, BX795, enhanced osimertinib response in resistant PDXs. **A)** pPDK1 (S241) expression in TC386-OSIR PDXs treated with BX 795, osimertinib and osimertinib + BX 795 B) Level of pPDK1 (S241) expression in osimertinib sensitive TC386 PDX, and osimertinib resistant TC386-OSIR PDX tissues comparing with H1975-OSIR/PDK-/- and H1975-OSIR/PDK++/++ cells, C) Osimertinib + BX795 treatment strategies in osimertinib resistant PDXs, D) Antitumor activity of osimertinib + BX795 combination on TC386-OSIR PDXs, E) Growth curves of TC386-OSIR PDXs bearing individual mice in different treatment groups.

### PDK1 knock-out alters activation of the AKT/mTOR pathway

Activation of the oncogenic PI3K/AKT/mTOR pathway mediates tumorigenesis and resistance to EGFR TKIs in NSCLC^16–19^. Since PDK1 represents a pivotal node in this important signaling axis, we analyzed the phosphorylation status of its major signaling effectors in NCI-H1975, NCI-H1975/OSIR, H1975-OsiR-PDK-/- and H1975-OsiR-PDK1++/++ clones. Figure 7A shows that AKT expression was similar in both H1975-OsiR-PDK1-/- and H1975-OsiR-PDK1++/++ clones. Osimertinib treatment had no effect on its level. In H1975-OsiR-PDK1++/++ clone, Osimertinib reduced AKT phosphorylation at threonine 308 (T308) residue, which is known to be the site activated by PDK1^23^. PDK1 knock-out had no effect on AKT (S473) phosphorylation, which can be catalyzed by multiple proteins but not PDK1. Although the level of AKT expression in OsiR-PDK1-/- cells was similar to that of H1975-OsiR-PDK1++/++, there was no detected phosphorylation of AKT (T308) in the former. The level of phosphorylation of AKT (S473) in H1975-OsiR-PDK^-/-^ was higher than that of H1975-OsiR-PDK1++/++ cells. Osimertinib had no effect on this activity. The level of mTOR expression was similar in H1975-OsiR-PDK^-/-^ and H1975-OsiR-PDK1++/++ clones, but its phosphorylation level was higher in the latter, and osimertinib had no effect on mTOR or pmTOR levels (Fig 7B). PTEN is a tumor suppressor that regulates the PI3K/AKT/PTEN pathway important in senescence and apoptosis^39^. Analysis of PTEN total expression showed that the expression of PTEN was downregulated in NCI-H1975/OSIR compared with NCI-H1975 (Fig 7C). Phosphorylation of PTEN was upregulated by osimertinib treatment only in NCI-H1975.

**Fig 7.**
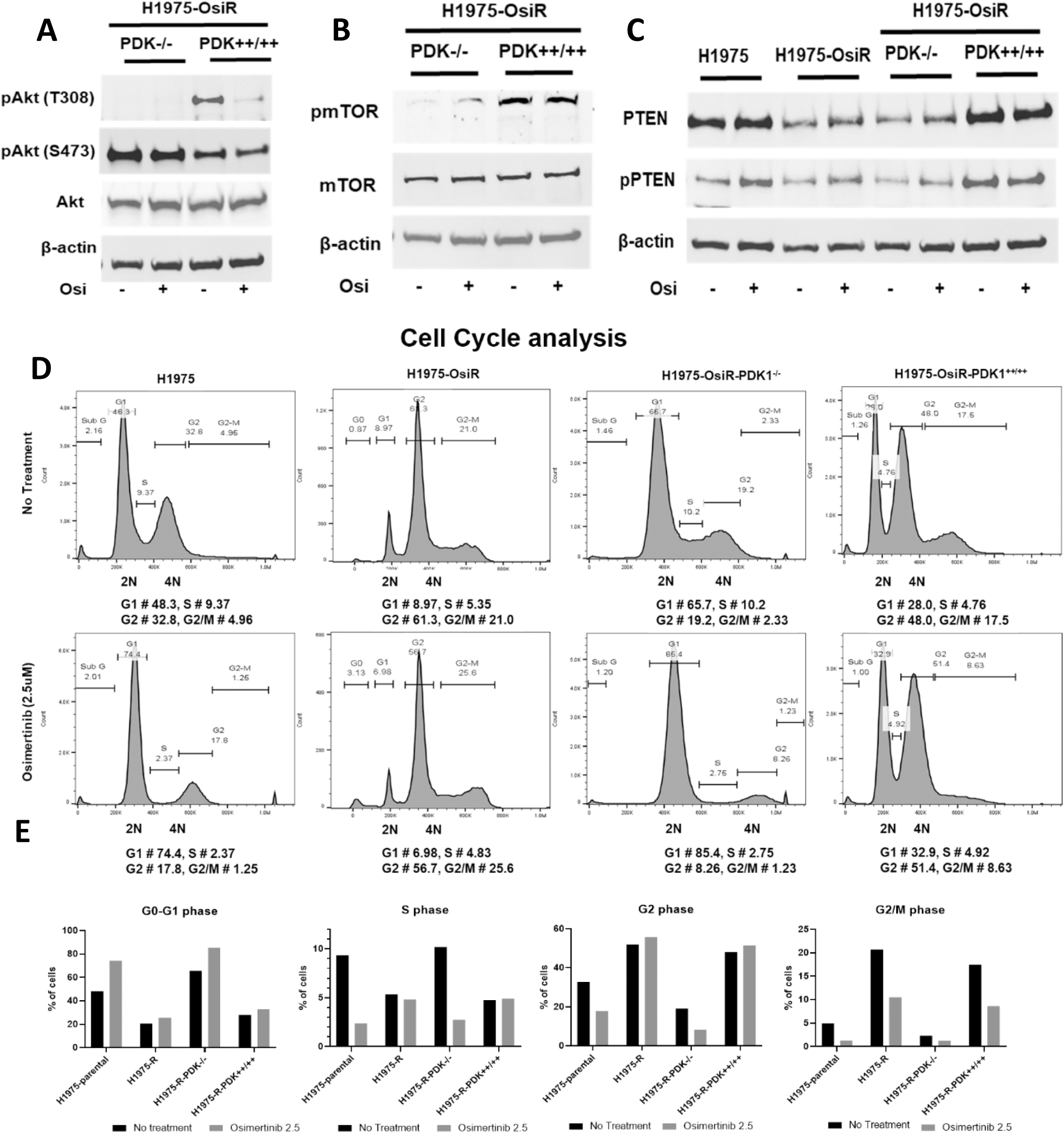
PDK1 knock-out dysregulates /Akt/mTOR signaling and promotes cell cycle arrest. A) Akt, B) mTOR and C) PTEN expression in H1975-OsiR-PDK1-/- and H1975-OsiR-PDK1++/++ cells and alteration by osimertinib treatment. C) PTEN expression status among H1975-parental, H1975-OsiR, H1975-OsiR-PDK1-/- and H1975-OsiR-PDK1++/++ cells. D) Cell cycle analysis of H1975-parental, H1975-OsiR, H1975-OsiR-PDK1-/- and H1975-OsiR-PDK1++/++ cells after osimertinib treatment. E) Quantitation of cells in difference phases and its alteration by osimertinib treatment.

### PDK1 knock-out promotes cell cycle arrest at G1

Cell cycle analysis showed that osimertinib promoted cell cycle arrest at G1 in NCI-H1975 sensitive cells. In contrast, a large number of NCI-H1975/OSIR cells accumulated at G2 and G2/M after osimertinib treatment (Fig 7D, E). The PDK1 knock out also promoted cell cycle arrest at the G1 phase, to the same extent as in NCI-H1975 sensitive cells. NCI-H1975 OSIR-PDK++/++ cells were not arrested at G1 post osimertinib treatment, which is similar to NCI-H1975/OSIR cells. These results suggest that PDK1 knock-out renders the resistant cells more sensitive to osimertinib through cell cycle arrest at G1.

### PDK1 knock-out inhibits pYAP expression and nuclear translocation of YAP phosphorylated at Y357

H1975-OsiR-PDK^-/-^ downregulated YAP and pYAP whereas NCI-H1975 OSIR-PDK++/++ had increased levels of YAP and pYAP expression (Fig 8A-B). In osimertinib sensitive cells, NCI-H1975, osimertinib treatment downregulated YAP and pYAP. However, no osimertinib effect on YAP was noted for NCI-H1975/OSIR cells (Fig 8A-B). NCI-H1975/OSIR and NCI-H1975 OSIR-PDK++/++ had higher levels of pYAP than H1975-OsiR-PDK^-/-^ . Phosphorylation of YAP at Y357 is activating and promotes translocation of YAP to the nucleus. An anti-pYAP^Y357^ antibody was used to localize YAP^Y357^ by immunofluorescence. Osimertinib treatment of H1975 osimertinib sensitive cells reduced nuclear localization of pYAP^Y357^. NCI-H1975/OSIR and NCI-H1975 OSIR-PDK++/++ cells showed a high level of nuclear localization of pYAP^Y357^. H1975-OsiR-PDK-/- cells showed significantly reduced nuclear localization of pYAP^Y357^ (Fig 8C-D). The level of YAP and pYAP was upregulated in osimertinib resistant xenograft tumors as well as in residual tumor tissues following osimertinib treatment (Fig 8E).

**Fig 8.**
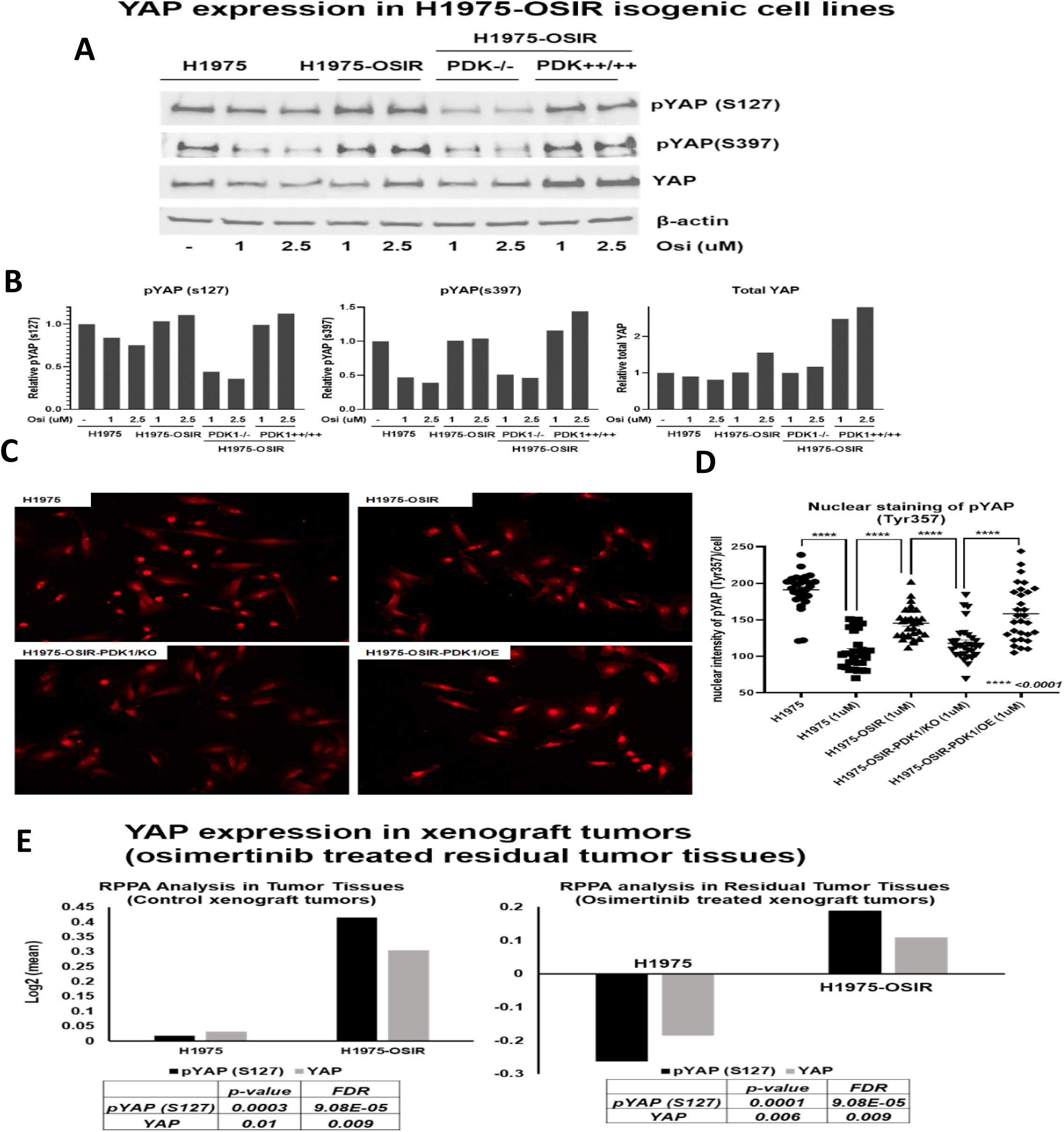
PDK1 knock-out inhibits YAP expression and nuclear translocation. A) Western blot shows expression of total YAP, pYAP (S127), pYAP (S397) in osimertinib sensitive H1975, resistant H1975-OSIR, PDK1-/- and PDK1++/++ cells, B) Quantitation of pYAP (S127), pYAP (S397) and total YAP in H1975 isogenic cell lines, C) Immunofluorescence images of nuclear translocation of YAP detected by immunostaining with pYAP (Tyr 357) antibody on osimertinib sensitive and resistant cell lines as well as PDK1 KO and OE cells, D) quantitative analysis of nuclear YAP signals in four H1975 isogenic cell lines, E) level of YAP and pYAP (S127) in osimertinib sensitive and resistant xenograft tumors (left panel) and osimertinib treated residual sensitive and resistant tumors (right panel).

### Immunohistochemical analysis shows an association between high PDK1 expression level and progressive disease in EGFR mutant NSCLC patients

Formalin-fixed paraffin-embedded (FFPE) tumor blocks from EGFR mutant NSCLC tumors obtained prior to initiation of treatment or after tumor progression during treatment were stained with anti-PDK1 antibody for IHC analysis. The highest levels of PDK1 expression were only observed in the progressive disease patients, suggesting they could be responsive to an osimertinib and PDK1 inhibitor combination (Supplemental Fig 5).

## Discussion

Responses to osimertinib and other TKIs are transient, and acquired resistance is inevitable. The majority of EGFR mutant NSCLC patients treated initially with osimertinib will eventually progress after only 19 months of treatment ^40^. Although acquired new mutations in EGFR account for some clinical acquired resistance to both osimertinib and other TKIs, the majority of resistant phenotypes cannot be explained by acquisition of these mutations. Delaying and treating tumors with acquired resistance requires an understanding of multiple complex resistance mechanisms mediated by alternative bypass pathways. In this study, we used the extensively molecularly characterized human NSCLC NCI-H1975 cell line which harbors two common EGFR point mutations, T790M and L858R, in exons 20 and 21 respectively, which is very sensitive to osimertinib, and its isogenic derivative osimertinib resistant clone as a model system, to first develop a relevant humanized mouse model that models osimertinib acquired resistance accurately, and second, decipher cellular and molecular mechanisms for its acquired resistance. The NCI-H1975 resistant clone, NCI-H1975/OSIR, is 100-200 fold more resistant to osimertinib. Whole exome sequencing eliminated the acquisition of new EGFR mutations or loss of T790M and L858R in the resistant clone, suggesting the existence of alternative mechanisms of resistance acquired during osimertinib treatment.

Osimertinib sensitive and resistant humanized xenografts had different responses to short and long-term treatment, with the latter having initially a slowing of growth followed by aggressive tumor regrowth. Sensitive tumors had very long regression, before tumors started to regrow slowly similar to responses seen in the clinic. When compared to current mouse models, our humanized mouse model replicates human EGFR mutant tumor growth physiology, pathology, immunology, and response to osimertinib treatment. In our recent published report, we showed that HLA matching between CD34 stem-cell donors and inoculated tumors or implanted PDXs, was not necessary for an antigen-specific antitumor response^31^. Using this model, we investigated differences between reconstituted human immune contextures in the TME of osimertinib sensitive and resistant xenografts in humanized mice. The results showed that osimertinib sensitive tumors had a higher number of M1 macrophages, dendritic cells, and infiltrating lymphocytes, all of which are strong immunostimulatory contributors to the TME. Osimertinib treatment enhanced their intratumor levels significantly in both sensitive and resistant tumors. On the other hand, the level of M2 macrophages was higher in resistant tumors, and osimertinib decreased their levels significantly. This immune cell population is a strong contributor to the immunosuppressive tumor microenvironment, and its preponderance is an obstacle for immunotherapy. Osimertinib significantly altered the ratio of M1 to M2, favoring the M1 population which has an immunostimulatory phenotype. Unexpectedly, the immunosuppressive level of CD33^+^ MDSC levels were slightly higher in sensitive compared to resistant tumors and were markedly decreased after osimertinib treatment.

Pairwise comparison analysis of protein expression profiles of residual tumors in osimertinib treated NSG mice, using reverse phase protein array (RPPA), between NCI-H1975 compared to NCI-H1975/OSIR, and NCI-H1975/OSIR osimertinib treated vs untreated groups, showed two distinct protein signatures. We found unexpectedly that both signatures were led by upregulation of PDK1, a master regulator the AGC kinase superfamily^22^. PDK1 regulates the oncogenic PI3k/AKT/mTOR pathway, which is involved in tumorigenesis and progression of NSCLC. Next, global and phospho-proteome based mass spectrometry (MS Spec) analyses between NCI-H1975 and NCI-H1975/OSIR clones did not find any detectable increase in PDK1 expression level but indicated a highly significant fourteen fold increase of phosphorylated PDK1 in the resistant clone. Pharmacological and genetic suppression of PDK1 sensitized NCI-H1975/OSIR cells to osimertinib. Cell survival and colony formation assays showed that NCI-H1975/OSIR clones treated with the specific PDK1 inhibitor, BX795, or PDK1 knock out by CRISPR/Cas9, recovered their sensitivity to osimertinib treatment. In addition, NCI-H1975/OSIR PDK1 knockout clone H1975-OsiR-PDK-/-, with restored overexpression of PDK1 was more resistant to osimertinib than its parental clone, validating the role of PDK1 in mediating osimertinib resistance in this model. Treatment with a combination of osimertinib and BX795 in a second model of acquired osimertinib resistance utilizing a PDX showed that the addition of BX795 to osimertinib resulted in synergistic tumor regression whereas BX795 treatment did not differ from untreated control growth and osimertinib slowed growth but did not cause regression in the drug resistant PDX. This PDX model with acquired osimertinib resistance without a T790M mutation replicates a common clinical scenario that suggests a combination of osimertinib with a PDK1 inhibitor may be effective after progression on first-line osimertinib.

PDK1 is a pivotal node in the oncogenic PI3K/AKT/mTOR pathway^20,21^, which mediates tumorigenesis and resistance to EGFR tyrosine kinase inhibitors in NSCLC^16–19^. Osimertinib had no effect on AKT expression levels, which was similar in both PDK1 knock out and overexpressing clones. In the latter, osimertinib reduced AKT phosphorylation at the threonine 308 (T308) residue, which is known to be the site activated by PDK1. The level of mTOR expression was also similar in PDK1 knock out and overexpressing clones, but its phosphorylation level was higher in the latter indicating that PDK1 knock out can downregulate mTOR activation. A previous study by our group implicated mTOR as a mediator of TKI resistance^41^. In this study three NSCLC cell lines became sensitive to erlotinib following treatment with the mTOR inhibitor rapamycin.

PDK1 is also a mediator of yes-associated protein (YAP) activation^42^. PI3K and PDK1 mediate YAP phosphorylation and nuclear accumulation, and thus it is the PI3K-PDK1 signal that links EGFR with the Hippo pathway. Phosphorylation of YAP at Y357 is activating, resulting in YAP nuclear translocation^43^. Overexpression of PDK1 significantly increased YAP Y357 nuclear translocation in H1975 cells when compared to the PDK1 knockout, H1975-OsiR-PDK-/-, implicating YAP as a downstream mediator of osimertinib acquired drug resistance. This is supported by studies implicating YAP activation in persister cells after EGFR TKI treatment^44^. We recently reported YAP-driven transcriptional adaptation as a functional mechanism of TKI drug tolerance^45^. In this study, we found in experiments in humanized mice that YAP reduced treatment sensitivity to osimertinib and enhanced an immunosuppressive tumor microenvironment supporting tumor growth. Thus PDK1 is a central upstream regulator of two critical drug resistance pathways: PI3K/AKT/mTOR and YAP. This suggests that drugs targeting PDK1 could be beneficial in delaying the onset of acquired drug resistance and treating acquired drug resistance at its onset.

Cell cycle analysis showed that both osimertinib treatment of sensitive cells and PDK1 knock out promoted cell cycle arrest at the G1 phase, whereas resistant and PDK1 overexpressors were not arrested at G1. PDK1 is known to have a critical role in cell proliferation and cell cycle progression^46^. Finally, we showed that high expression of PDK1 was associated with progressive disease in EGFR mutant NSCLC patients, as shown by immunohistochemistry (IHC), suggesting they could be responsive to osimertinib and PDK1 inhibitor combination therapy.

In conclusion, we presented multiple lines of evidence for PDK1 as a driver of osimertinib acquired resistance in T790M/L858R mutant NSCLC using the most relevant preclinical mouse models, capable of modeling osimertinib acquired resistance, and interrogating its interaction with the microenvironments of sensitive and resistant tumors. We showed that pharmacological and genetic targeting of PDK1 could restore osimertinib responsiveness in cell lines and PDXs with acquired osimertinib resistance thus providing support for clinical translation.

## References

1. Gelatti, A. C. Z., Drilon, A. & Santini, F. C. Optimizing the sequencing of tyrosine kinase inhibitors (TKIs) in epidermal growth factor receptor (EGFR) mutation-positive non-small cell lung cancer (NSCLC). Lung Cancer 137, 113–122, doi:10.1016/j.lungcan.2019.09.017 (2019).

2. Kalemkerian, G. P. et al. Molecular Testing Guideline for the Selection of Patients With Lung Cancer for Treatment With Targeted Tyrosine Kinase Inhibitors: American Society of Clinical Oncology Endorsement of the College of American Pathologists/International Association for the Study of Lung Cancer/Association for Molecular Pathology Clinical Practice Guideline Update. J. Clin. Oncol. 36, 911–919, doi:10.1200/JCO.2017.76.7293 (2018).

3. Janne, P. A., et al. AZD9291 in EGFR inhibitor-resistant non-small-cell lung cancer. N. Engl. J. Med. 372, 1689–1699 (2015).

4. Govindan, R. Overcoming resistance to targeted therapy for lung cancer. N. Engl. J. Med. 372, 1760–1761, doi:10.1056/NEJMe1500181 (2015).

5. Yun, C. H. et al. The T790M mutation in EGFR kinase causes drug resistance by increasing the affinity for ATP. Proc. Natl. Acad. Sci. U. S. A. 105, 2070–2075, doi:10.1073/pnas.0709662105 (2008).

6. Sos, M. L. et al. Chemogenomic profiling provides insights into the limited activity of irreversible EGFR Inhibitors in tumor cells expressing the T790M EGFR resistance mutation. Cancer Res. 70, 868–874, doi:10.1158/0008-5472.CAN-09-3106 (2010).

7. Ramalingam, S. S. et al. Osimertinib As First-Line Treatment of EGFR Mutation-Positive Advanced Non-Small-Cell Lung Cancer. J. Clin. Oncol. 36, 841–849, doi:10.1200/JCO.2017.74.7576 (2018).

8. Ettinger, D. S. et al. Non-Small Cell Lung Cancer, Version 5.2017, NCCN Clinical Practice Guidelines in Oncology. J. Natl. Compr. Canc. Netw. 15, 504–535, doi:10.6004/jnccn.2017.0050 (2017).

9. Yang, J. C. et al. Osimertinib in Pretreated T790M-Positive Advanced Non-Small-Cell Lung Cancer: AURA Study Phase II Extension Component. J. Clin. Oncol. 35, 1288–1296, doi:10.1200/JCO.2016.70.3223 (2017).

10. Goss, G. et al. Osimertinib for pretreated *EGFR* Thr790Met-positive advanced non-small-cell lung cancer (AURA2): a multicentre, open-label, single-arm, phase 2 study. The Lancet Oncology 17, 1643–1652, doi:10.1016/S1470-2045(16)30508-3 (2016).

11. Wu, Y. L. et al. CNS Efficacy of Osimertinib in Patients With T790M-Positive Advanced Non-Small-Cell Lung Cancer: Data From a Randomized Phase III Trial (AURA3). J. Clin. Oncol. 36, 2702–2709, doi:10.1200/JCO.2018.77.9363 (2018).

12. Lim, S. M., Syn, N. L., Cho, B. C. & Soo, R. A. Acquired resistance to EGFR targeted therapy in non-small cell lung cancer: Mechanisms and therapeutic strategies. Cancer Treat. Rev. 65, 1–10, doi:10.1016/j.ctrv.2018.02.006 (2018).

13. Papadimitrakopoulou, V. A. et al. Analysis of resistance mechanisms to osimertinib in patients with EGFR T790M advanced NSCLC from the AURA3 study. Ann. Oncol. 29, 741–741 (2018).

14. Leonetti, A. et al. Resistance mechanisms to osimertinib in EGFR-mutated non-small cell lung cancer. Br. J. Cancer 121, 725–737, doi:10.1038/s41416-019-0573-8 (2019).

15. Oxnard, G. R. et al. Assessment of Resistance Mechanisms and Clinical Implications in Patients With EGFR T790M-Positive Lung Cancer and Acquired Resistance to Osimertinib. JAMA oncology 4, 1527–1534, doi:10.1001/jamaoncol.2018.2969 (2018).

16. Fumarola, C., Bonelli, M. A., Petronini, P. G. & Alfieri, R. R. Targeting PI3K/AKT/mTOR pathway in non small cell lung cancer. Biochem. Pharmacol. 90, 197–207, doi:10.1016/j.bcp.2014.05.011 (2014).

17. Heavey, S., O’Byrne, K. J. & Gately, K. Strategies for co-targeting the PI3K/AKT/mTOR pathway in NSCLC. Cancer Treat. Rev. 40, 445–456, doi:10.1016/j.ctrv.2013.08.006 (2014).

18. Tan, A. C. Targeting the PI3K/Akt/mTOR pathway in non-small cell lung cancer (NSCLC). Thorac Cancer 11, 511–518, doi:10.1111/1759-7714.13328 (2020).

19. Iksen, Pothongsrisit, S. & Pongrakhananon, V. Targeting the PI3K/AKT/mTOR Signaling Pathway in Lung Cancer: An Update Regarding Potential Drugs and Natural Products. Molecules 26, doi:10.3390/molecules26134100 (2021).

20. Lien, E. C., Dibble, C. C. & Toker, A. PI3K signaling in cancer: beyond AKT. Curr. Opin. Cell Biol. 45, 62–71, doi:10.1016/j.ceb.2017.02.007 (2017).

21. Faes, S. & Dormond, O. PI3K and AKT: Unfaithful Partners in Cancer. Int J Mol Sci 16, 21138–21152, doi:10.3390/ijms160921138 (2015).

22. Arencibia, J. M., Pastor-Flores, D., Bauer, A. F., Schulze, J. O. & Biondi, R. M. AGC protein kinases: from structural mechanism of regulation to allosteric drug development for the treatment of human diseases. Biochim. Biophys. Acta 1834, 1302–1321, doi:10.1016/j.bbapap.2013.03.010 (2013).

23. Scheid, M. P., Parsons, M. & Woodgett, J. R. Phosphoinositide-dependent phosphorylation of PDK1 regulates nuclear translocation. Mol. Cell. Biol. 25, 2347–2363, doi:10.1128/MCB.25.6.2347-2363.2005 (2005).

24. Hossen, M. J. et al. PDK1 disruptors and modulators: a patent review. Expert Opin. Ther. Pat. 25, 513–537, doi:10.1517/13543776.2015.1014801 (2015).

25. Hann, S. S., Zheng, F. & Zhao, S. Targeting 3-phosphoinositide-dependent protein kinase 1 by N-acetyl-cysteine through activation of peroxisome proliferators activated receptor alpha in human lung cancer cells, the role of p53 and p65. J. Exp. Clin. Cancer Res. 32, 43, doi:10.1186/1756-9966-32-43 (2013).

26. Ferro, R. & Falasca, M. Emerging role of the KRAS-PDK1 axis in pancreatic cancer. World J. Gastroenterol. 20, 10752–10757, doi:10.3748/wjg.v20.i31.10752 (2014).

27. Caron, R. W. et al. Activated forms of H-RAS and K-RAS differentially regulate membrane association of PI3K, PDK-1, and AKT and the effect of therapeutic kinase inhibitors on cell survival. Mol. Cancer Ther. 4, 257–270 (2005).

28. Lee, K. Y., D’Acquisto, F., Hayden, M. S., Shim, J. H. & Ghosh, S. PDK1 nucleates T cell receptor-induced signaling complex for NF-kappaB activation. Science 308, 114–118, doi:10.1126/science.1107107 (2005).

29. Yagi, M. et al. PDK1 regulates chemotaxis in human neutrophils. J. Dent. Res. 88, 1119–1124, doi:10.1177/0022034509349402 (2009).

30. Han, L. et al. Prognostic potential of microRNA-138 and its target mRNA PDK1 in sera for patients with non-small cell lung cancer. Med. Oncol. 31, 129, doi:10.1007/s12032-014-0129-y (2014).

31. Meraz, I. M. et al. An Improved Patient-Derived Xenograft Humanized Mouse Model for Evaluation of Lung Cancer Immune Responses. Cancer Immunol Res 7, 1267–1279, doi:10.1158/2326-6066.CIR-18-0874 (2019).

32. Andrews, S. FastQC: a quality control tool for high throughput sequence data. (2010).

33. Li, H. & Durbin, R. Fast and accurate short read alignment with Burrows-Wheeler transform. Bioinformatics 25, 1754–1760, doi:10.1093/bioinformatics/btp324 (2009).

34. McKenna, A. et al. The Genome Analysis Toolkit: a MapReduce framework for analyzing next-generation DNA sequencing data. Genome Res. 20, 1297–1303, doi:10.1101/gr.107524.110 (2010).

35. Wang, K., Li, M. & Hakonarson, H. ANNOVAR: functional annotation of genetic variants from high-throughput sequencing data. Nucleic Acids Res. 38, e164, doi:10.1093/nar/gkq603 (2010).

36. Hu, J. et al. Non-parametric quantification of protein lysate arrays. Bioinformatics 23, 1986–1994, doi:10.1093/bioinformatics/btm283 (2007).

37. Berenbaum, M. C. What is synergy? [published erratum appears in Pharmacol Rev 1990 Sep;41(3):422]. Pharmacol. Rev. 41, 93–141 (1989).

38. Huang, L. et al. CombPDX: a unified statistical framework for evaluating drug synergism in patient-derived xenografts. bioRxiv, 2021.2010.2019.464994, doi:10.1101/2021.10.19.464994 (2021).

39. Papa, A. & Pandolfi, P. P. The PTEN-PI3K Axis in Cancer. Biomolecules 9, doi:ARTN 153 10.3390/biom9040153 (2019).

40. Soria, J. C. et al. Osimertinib in Untreated EGFR-Mutated Advanced Non-Small-Cell Lung Cancer. N. Engl. J. Med. 378, 113–125, doi:10.1056/NEJMoa1713137 (2018).

41. Dai, B. et al. Exogenous restoration of TUSC2 expression induces responsiveness to Erlotinib in Wildtype Epidermal Growth Factor Receptor (EGFR) lung cancer cells through context specific pathways resulting in enhanced therapeutic efficacy. PLoS One 10, e0123967 (2015).

42. Emmanouilidi, A. & Falasca, M. Targeting PDK1 for Chemosensitization of Cancer Cells. Cancers (Basel) 9, doi:10.3390/cancers9100140 (2017).

43. Li, B. et al. c-Abl regulates YAPY357 phosphorylation to activate endothelial atherogenic responses to disturbed flow. J. Clin. Invest. 129, 1167–1179, doi:10.1172/JCI122440 (2019).

44. Kurppa, K. J. et al. Treatment-Induced Tumor Dormancy through YAP-Mediated Transcriptional Reprogramming of the Apoptotic Pathway. Cancer Cell 37, 104-+, doi:10.1016/j.ccell.2019.12.006 (2020).

45. Haderk, F. et al. A focal adhesion kinase-YAP signaling axis drives drug tolerant persister cells and residual disease in lung cancer. bioRxiv, 2021.2010.2023.465573, doi:10.1101/2021.10.23.465573 (2021).

46. Nakamura, K. et al. PDK1 regulates cell proliferation and cell cycle progression through control of cyclin D1 and p27(Kip1) expression. J. Biol. Chem. 283, 17702–17711, doi:10.1074/jbc.M802589200 (2008).

